# The ER-phagy receptor FAM134B is targeted by *Salmonella* Typhimurium to promote infection

**DOI:** 10.1101/2024.03.28.587227

**Authors:** Damián Gatica, Reham M. Alsaadi, Rayan El Hamra, Rudolf Mueller, Makoto Miyazaki, Subash Sad, Ryan C. Russell

**Affiliations:** Department of Cellular and Molecular Medicine, University of Ottawa, Ottawa, ON, Canada; Department of Biochemistry, Microbiology and Immunology, University of Ottawa, Ottawa, ON, Canada; Department of Pathology and Laboratory Medicine, Faculty of Medicine, University of Ottawa, Ottawa, ON, Canada; Division of Renal Diseases and Hypertension, Department of Medicine, University of Colorado Denver, Aurora, CO, USA; Ottawa Institute of Systems Biology, University of Ottawa, Canada; University of Ottawa Centre for Infection, Immunity and Inflammation, Ottawa, ON, Canada

**Keywords:** Xenophagy, Autophagy, Endoplasmic Reticulum, Bacteria, RETREG1

## Abstract

Macroautophagy/autophagy is a key catabolic-recycling pathway that can selectively target damaged organelles or invading pathogens for degradation. The selective autophagic degradation of the endoplasmic reticulum (hereafter referred to as ER-phagy) is a homeostatic mechanism, controlling ER size, the removal of misfolded protein aggregates, and organelle damage. ER-phagy is also stimulated by pathogen infection. However, the link between ER-phagy and bacterial infection remains poorly understood, as are the mechanisms evolved by pathogens to escape the effects of ER-phagy. Here, we show that *Salmonella enterica* serovar Typhimurium inhibits ER-phagy by targeting the ER-phagy receptor FAM134B, leading to a pronounced increase in *Salmonella* viability after invasion. *Salmonella* prevents FAM134B oligomerization, which is required for efficient ER-phagy. FAM134B knock-out raises intracellular *Salmonella* number, while FAM134B activation reduces *Salmonella* burden. Additionally, we found that *Salmonella* targets FAM134B through the bacterial effector SopF to enhance intracellular survival through ER-phagy inhibition. Furthermore, FAM134B knock-out mice infected with *Salmonella* presented severe intestinal damage and increased bacterial burden. These results provide new mechanistic insight into the interplay between ER-phagy and bacterial infection, highlighting a key role for FAM134B in innate immunity.

## INTRODUCTION

Macroautophagy (hereafter autophagy) is an intracellular catabolic-recycling pathway that promotes survival in response to a diverse range of stress conditions including nutrient starvation or invasive pathogens^1^. Autophagic cargo can include proteins, lipids, damaged organelles, or even intracellular pathogens. These cytoplasmic components are sequestered inside double-membrane vesicles called autophagosomes. Fully formed autophagosomes then fuse with lysosomes leading to the degradation of the sequestered cargo by resident hydrolytic enzymes. The basic macromolecules obtained from cargo degradation are subsequently transported back to the cytoplasm for reutilization. Autophagy levels are regulated by several Atg (autophagy-related) proteins, involved in all steps of the autophagic process from initiation to lysosomal fusion^2^. Autophagosome cargo selection is often tightly regulated, specifically targeting damaged cellular components or invasive intracellular pathogens for degradation^3^. Selective autophagy is achieved by autophagy receptors that link the cargo targeted for degradation with the growing autophagosomal membrane. Autophagy receptors usually contain LC3-interacting regions (LIR), an evolutionary conserved sequence that binds to members of the Atg8/LC3/GABARAP family^4^. During autophagy initiation LC3 is lipidated and covalently bound to the growing autophagosomal membrane. Thus, by interacting with LC3, autophagy receptors selectively tether cargo to the sequestering autophagosome^3,4^. The selective autophagic degradation of the endoplasmic reticulum (ER), termed ER-phagy, has been shown to be necessary to mitigate ER stress and maintain ER homeostasis. ER-phagy controls ER size and morphology, as well as inducing the degradation of misfolded protein aggregates that can be toxic for the cell^3,5^. Several ER-phagy cargo receptors have been described, including FAM134B^6^ and TEX264^7,8^, among others^5^. Recent reports have begun to describe the different molecular mechanisms by which ER-phagy receptors are regulated in the promotion of ER-phagy^9–11^. The first ER-phagy receptor identified, FAM134B, is an ER transmembrane protein containing a reticulon-homology domain that allows it to sense and induce ER membrane curvature and budding through protein clustering^12,13^. FAM134B activity requires its oligomerization, which is highly regulated by different post-translational modifications, and is a requisite step in FAM134B-driven ER-phagy^9,10,14^. Interestingly, FAM134B has also been involved in the cellular response against viral infection. FAM134B-dependent ER-phagy has been shown to limit SARS-COV-2, Ebola, Zika and dengue virus replication^15–17^. Moreover, multiple pathogens have developed strategies to inhibit ER-phagy by specifically hijacking or cleaving FAM134B^16,17^. Targeting FAM134B leads to ER remodeling, which is thought to benefit invading viruses, creating a favorable environment for replication^17,18^. However, the links between ER-phagy and infection; and the mechanisms pathogens have evolved to evade the effects of ER-phagy remain poorly understood. In this study, we identified a novel mechanism of bacterial-mediated inhibition of ER-phagy. Specifically, we found that *Salmonella enterica* serovar Typhimurium (*Salmonella*), a major cause of foodborne infections worldwide^19^, inhibits ER-phagy by specifically targeting the ER-phagy receptor FAM134B, leading to a pronounced increase in *Salmonella* viability after invasion. We show that *Salmonella* infection prevents FAM134B oligomerization, which is required for efficient ER-phagy. Conversely, *Salmonella* -mediated ER-phagy blockage could be bypassed by promoting FAM134B oligomerization, which recovered ER-phagy levels. We provide evidence that *Salmonella* targets FAM134B through the bacterial effector SopF, preventing oligomerization and ER-phagy activation. Furthermore*, in vivo* analysis of FAM134B knock-out mice infected with *Salmonella* revealed intestinal damage and increased bacterial levels in the spleen, colon and feces. Our results provide new mechanistic insight into the interplay between ER-phagy and bacterial infection, highlighting a key role for FAM134B in innate immunity.

## RESULTS

### *Salmonella* Typhimurium infection blocks ER-phagy

To investigate the possible impact of *Salmonella* infection on ER-phagy, we generated a HEK293 cell line stably expressing a previously published doxycycline-inducible ER-phagy reporter containing an ER signal sequence, followed by the fluorescent proteins RFP and GFP, and the ER retention sequence KDEL (ss-RFP-GFP-KDEL)^7^. Upon doxycycline treatment and ER-phagy induction, the fluorescent ER-associated reporter is sequestered inside autophagosomes and cleaved upon lysosome fusion. Unlike GFP, RFP is relatively resistant to lysosomal pH and hydrolases^20^. As such, a 25-KDa fragment corresponding to RFP can be detected by western blot upon activation of ER-phagy. Similarly, the same reporter can be used to measure ER-phagy by fluorescent microscopy due to the quenching of GFP fluorescence at lysosomal acidic pH (Fig. 1A). As a result, ER-phagy also can be monitored as an increase in RFP positive structures (lysosome associated ER-phagy probe) compared to the dual positive probe signal. We tested the reporter by activating ER-phagy through amino acid starvation or Torin-1 (a potent autophagy inducer) treatment. As a negative control, we confirmed that ss-RFP-GFP-KDEL did not get cleaved in autophagy-deficient *FIP200* knock-out (KO) cells (Supp Fig. 1A).

**Fig. 1.**
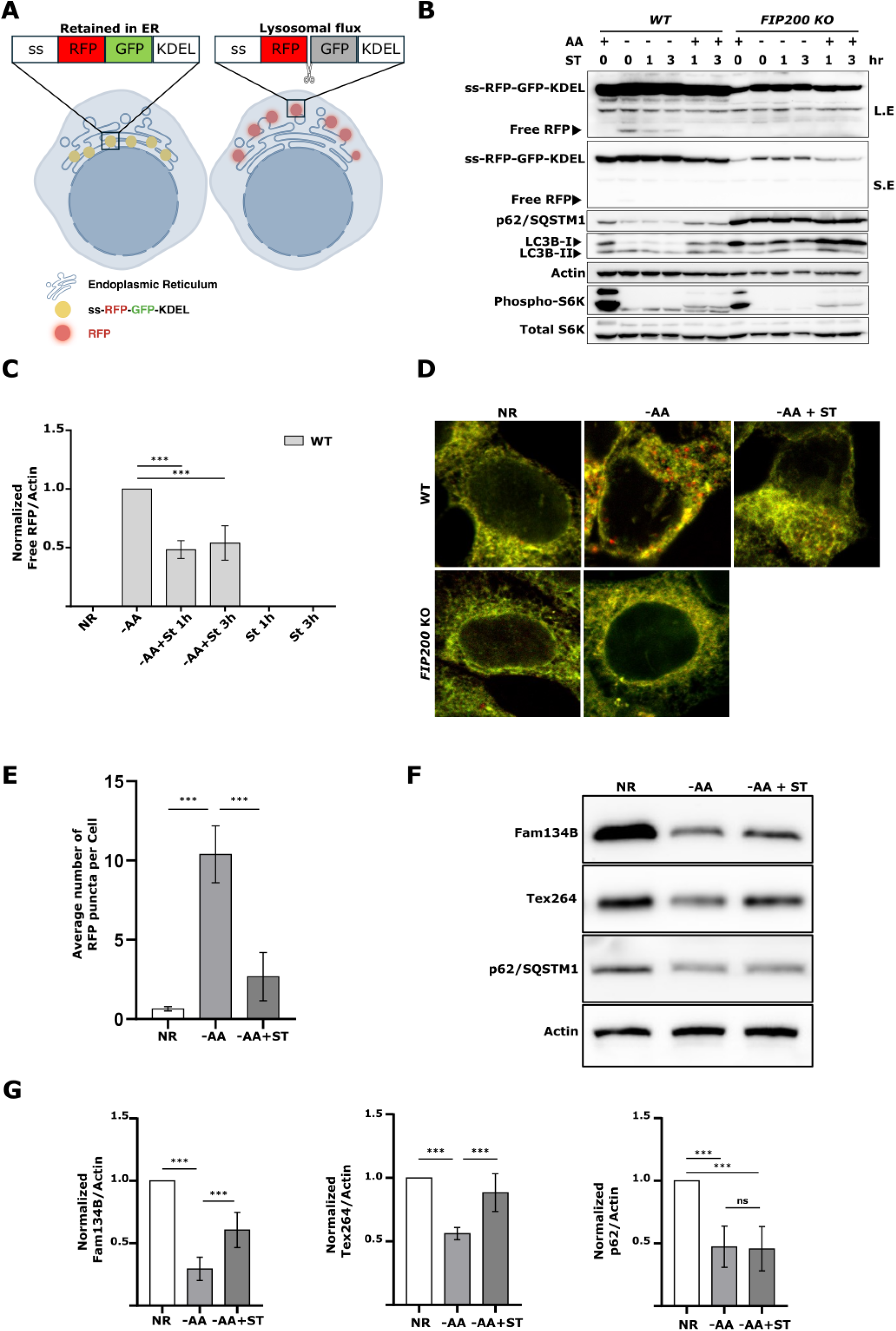
*Salmonella* infection blocks ER-phagy. **A)** Schematic for the ER-phagy probe ss-RFP-GFP-KDEL. During ER-phagy ss-RFP-GFP-KDEL is processed by lysosomal hydrolases to generate a free RFP fragment. Lysosomal acidity quenches the GFP signal. ss, signal sequence (Created with BioRender.com). **B)** WT and *FIP200* KO HEK293 cells stably expressing the ss-RFP-GFP-KDEL reporter were starved for AA for 6h; starved for 3h followed by ST infection for 1 or 3h in AA starvation media; or infected in nutrient rich media with ST for 1 or 3h. ER-phagy was measured by ss-RFP-GFP-KDEL processing. Non-selective autophagy activity was measured by LC3B lipidation and p62/SQSTM1 degradation. Phospho- and total S6K levels were determined to ascertain AA starvation. Actin was used as a loading control. AA, amino acids; ST, *Salmonella* Typhimurium; S.E, Short Exposure; L.E, Long Exposure. **C)** Normalized Free RFP-Actin ratio from WT cells in A). Error bars indicate the standard deviation of 3 independent experiments. ANOVA, ***P <0.001. NR, Nutrient Rich. D) Cells from A) were either AA starved for 6h or starved for 3h followed by ST infection for 3h in AA starvation media, fixed and image by confocal microscopy. **E)** Average number of RFP puncta per cell from D) were quantified. ANOVA, ***P <0.001. **F)** WT cells were AA starved for 2h or starved for 2h in the presence of ST. Fam134B, Tex264, p62/SQSTM and Actin levels were determined by western blot. **G)** Normalized FAM134B-Actin, TEX264-Actin and p62/SQSTM ratios from F) were quantified. ns, no significance. Error bars indicate the standard deviation of at least 5 independent experiments. ANOVA, ***P <0.001.

Amino acid starvation for 6 hr was sufficient to induce ER-phagy. To determine if *Salmonella* infection impacted ER-phagy, we added *Salmonella* to the minus amino acid media for the final 1 or 3 hrs of starvation. Both 1 and 3 hr of *Salmonella* infection significantly reduced the production of the free RFP fragment when compared to 6 hr of amino acid starvation in the absence of *Salmonella* (Fig. 1B and 1C), indicating inhibition of ER-phagy upon infection. Interestingly, *Salmonella* infection failed to block non-selective autophagy as shown by the degradation of p62/SQSTM1, a cargo receptor regularly used as an index for general autophagic degradation and LC3B-II production, the faster electrophoretic form of LC3B that is produce during autophagy initiation by LC3B lipidation^21,22^ (Fig. 1B). Similarly, CYTO-ID, a commercial autophagy detection kit, confirmed that non-selective autophagy was not affected by *Salmonella* infection during starvation, suggesting *Salmonella* specifically targets ER-phagy (Supp Fig. 1B). Importantly, Bafilomycin A1 (BAFA1) treatment, a potent inhibitor of autophagosome maturation and cargo degradation, completely blocked free RFP production when cells were starved in the presence of *Salmonella*, indicating that *Salmonella* did not increase the turnover rate of autolysosomes (Supp Fig. 1C). Utilizing the ability of the probe to visualize ER-phagy by fluorescence microscopy, we found that *Salmonella* infection significantly decreased ER-phagy induction (RFP puncta formation) compared to non-infected cells (Fig. 1D and 1E), which was consistent with our western blot analysis. We next measured the protein levels of endogenous ER-phagy cargo receptors known to be involved in starvation-induced ER-phagy, namely *TEX264* and *FAM134B*. Because these receptors link the ER to the autophagosome membrane they are ultimately degraded making their protein levels inversely correlated with ER-phagy levels^6,7^. We measured the endogenous level of each receptor during ER-phagy inducing conditions in the absence and presence of *Salmonella* and found both TEX264 and FAM134B levels to be significantly higher upon infection, indicating that their degradation is blocked when *Salmonella* infected the cells (Fig. 1F and 1G). Conversely, p62/SQSTM1 degradation was similar between infected and non-infected samples. Collectively, these findings indicate that *Salmonella* infection specifically inhibits ER-phagy, but not non-selective autophagy.

### FAM134B is targeted by *Salmonella* to block ER-phagy

The ER-phagy receptor proteins FAM134B and TEX264 have been previously reported to be targeted by invasive pathogens to disrupt ER morphology and promote infection^18^. To investigate this possibility, we generated CRISPR KO cell lines of *FAM134B* and *TEX264* (Supp Fig. 2A). As expected, deletion of *FAM134B* and *TEX264* severely decreased ER-phagy (Fig 2A), which is in line with prior reports^7^. Consistent with our previous findings, WT cells infected with *Salmonella* showed a significant decrease in the levels of free RFP compared to uninfected cells during ER-phagy inducing conditions. However, *Salmonella* infection failed to show a significant difference in ER-phagy levels between infected and uninfected *FAM134B* KO cells. In contrast, *Salmonella* was still capable of repressing ER-phagy in *TEX264* KO cells (Fig. 2A and 2B). Immunofluorescence microscopy using the same experimental setup showed a dramatic decrease in RFP puncta formation when ER-phagy was induced in *Salmonella* -infected WT and *TEX264* KO cells. However, RFP puncta formation in *FAM134B* KO cells infected with *Salmonella* during ER-phagy induction failed to decrease to the same extent, displaying significant differences (Fig. 2C and 2D). These results suggest that *Salmonella* infection primarily targets FAM134B to inhibit ER-phagy.

**Fig. 2.**
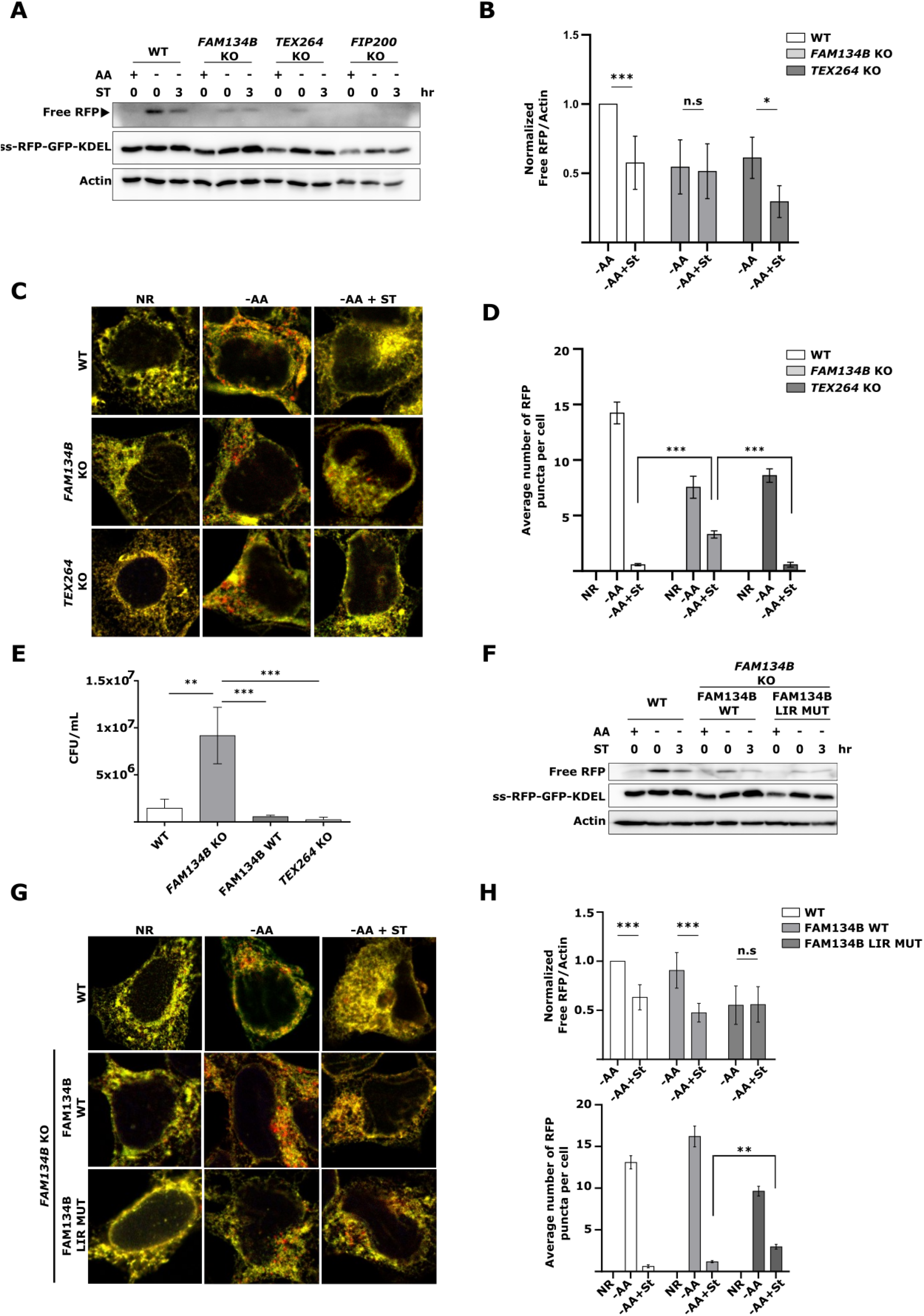
FAM134B is targeted by *Salmonella* to block ER-phagy. **A)** WT, *FAM134B* KO, *TEX264* KO and *FIP200* KO HEK293 cells stably expressing the ss-RFP-GFP-KDEL were either kept in nutrient rich media, starved for AA for 6h or starved for 3h followed by ST infection for 3h in AA starvation media. ER-phagy was measured by ss-RFP-GFP-KDEL processing. Actin was used as a loading control. AA, amino acids; ST, *Salmonella* Typhimurium; SE, Short Exposure; LE, Long Exposure. **B)** Normalized Free RFP-Actin ratio from cells in A). Error bars indicate the standard deviation of 7 independent experiments. ANOVA, *P <0.05; ***P <0.001; n.s, no significance. **C)** Cells from A) were either AA starved for 6h, or AA starved for 3h, followed by ST infection for 3h in starvation media, fixed and imaged by confocal microscopy. NR, Nutrient Rich. **D)** Average number of RFP puncta per cell from C) were quantified. ANOVA, ***P <0.001. **E)** WT, *FAM134B* KO, *TEX264* KO and *FAM134B* KO cells transfected with WT FAM134B were infected with *Salmonella*. Bacterial content was determined through a colony-forming unit (CFU). Error bars indicate standard deviation. ANOVA, **P <0.01; ***P <0.001. **F)** WT and *FAM134B* KO cells transfected with either FAM134B WT, FAM134B LIR mutant or mock were kept in nutrient rich media, starved for AA for 6h or starved for 3h followed by ST infection for 3h in AA starvation media. ER-phagy was measured by ss-RFP-GFP-KDEL processing. Actin was used as a loading control. **G)** Cells from F) were either AA starved for 6h, or AA starved for 3h, followed by ST infection for 3h in starvation media, fixed and imaged by confocal microscopy. **H)** Normalized Free RFP-Actin ratio from cells in F) and average number of RFP puncta per cell from G) were quantified. Error bars indicate the standard deviation of 5 independent experiments. Student’s t-test, **P <0.01; ***P <0.001; n.s, no significance.

Next, we sought to determine if *Salmonella-* mediated ER-phagy inhibition impacted intracellular bacterial viability. To this end, we performed a colony-forming unit (CFU) assay in WT, *FAM134B* KO, *TEX264* KO and *FAM134B* KO cells transfected with FAM134B WT. Analysis of *Salmonella* viability at 4 hours post-infection revealed that *FAM134B* KO cells had significantly higher number of viable internalized bacteria, suggesting a defect in the clearance of *Salmonella* when compared to WT and *TEX264* KO cells. Furthermore, *FAM134B* KO cells reconstituted with *FAM134B* recovered similar levels of *Salmonella* viability as parental cells (Fig. 2E), indicating FAM134B, but not TEX264, is important for *Salmonella* clearance.

ER-phagy receptors couple the ER to the autophagosomal membrane through LIRs. We next sought to determine if the impact of FAM134B on *Salmonella* growth was due to ER-phagy induction or another uncharacterized autophagy-independent function. To this end, we transfected *FAM134B* KO cells with either FAM134B WT or FAM134B LIR mutant and performed the ER-phagy reporter RFP processing assay. Intriguingly, we observed no significant difference between infected and uninfected ER-phagy induced cells when the FAM134B LIR mutant was expressed. Conversely, cells expressing FAM134B WT showed a significant decreased in RFP production when infected with *Salmonella* (Fig. 2F and 2H). Consistent with our western blot analysis, we also observed that the decrease in the number of RFP puncta in the FAM134B LIR mutant cells triggered by *Salmonella* infection was not as pronounced as the one in infected cells expressing FAM134B WT (Fig. 2G and 2H). When we quantified *Salmonella* viability by CFU, results showed that the expression of the FAM134B LIR mutation resulted in a significant increase in the number of *Salmonella* compared to cells expressing FAM134B WT (Supp Fig. 2B). Altogether, these findings suggest that FAM134B plays a crucial role in improving *Salmonella* clearance and that this role is connected to FAM134B ability to induce ER-phagy.

### FAM134B oligomerization is hindered by *Salmonella* infection

FAM134B promotes ER membrane scission and ER-phagy through its ability to oligomerize^9^. Therefore, we hypothesized that *Salmonella* infection might prevent ER-phagy by repressing FAM134B oligomerization. To test this hypothesis, FAM134B-FLAG was immunoprecipitated (IP) after 3 hr of ER-phagy induction in the presence and absence of *Salmonella* infection. Consistent with previous reports, FAM134B oligomers were relatively resistant to denatured conditions and are observed by western blot in a distinct, slow migrating band^9,14^. Notably, *Salmonella* infection significantly reduced the formation of FAM134B oligomers (Fig. 3A). Furthermore, amino acid starvation-induced FAM134B puncta formation was dramatically reduced by *Salmonella* infection, revealing *Salmonella* blocks ER membrane scission, which is driven by FAM134B oligomerization (Fig. 3B). These results indicate *Salmonella* may block ER-phagy by preventing the oligomerization of FAM134B. If *Salmonella* represses FAM134B by preventing oligomerization, then forcing FAM134B oligomerization would be predicted to bypass *Salmonella* -mediated ER-phagy reduction. To this end, we generated a FAM134B G216R mutant, a naturally occurring mutation that resides in FAM134B reticulon-homology domain and has been described to dramatically enhance FAM134B oligomerization^9,23^. We transfected either FAM134B WT or FAM134B G216R in *FAM134B* KO cells and measured ER-phagy through the RFP processing assay. As expected, cells transfected with FAM134B WT exhibited reduced ER-phagy under stimulated conditions when infected with *Salmonella* (Fig. 3C and 3D). However, cells transfected with FAM134B G216R showed no significant difference in ER-phagy rates in the presence or absence of *Salmonella*, indicating that *Salmonella* inhibits ER-phagy upstream, or at the level, of FAM134B activation. Analysis of ER-phagy by immunofluorescence under the same conditions, showed that FAM134B G216R largely prevented ER-phagy repression by *Salmonella,* compared to FAM134B WT (Fig. 3E and 3F). Together, these experiments support a model in which *Salmonella* regulates ER-phagy through repression of FAM134B activity.

**Fig. 3.**
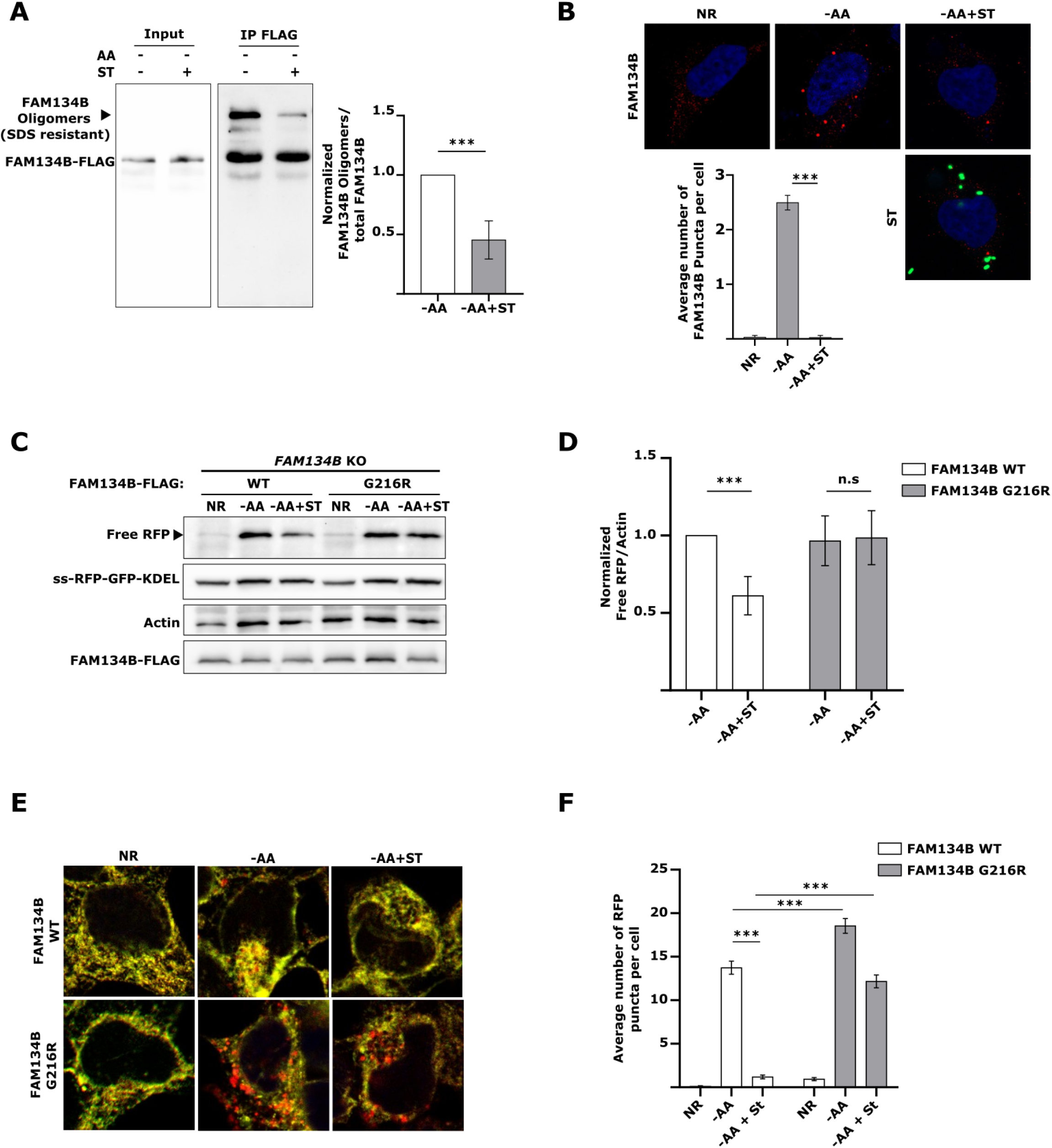
FAM134B oligomerization is hindered by *Salmonella* infection. **A)** *FAM134B* KO HEK293 cells transfected with FAM134B-FLAG were starved for AA for 3h in the absence or presence of ST. FLAG was immunoprecipitated. Error bars indicate the standard deviation of 5 independent experiments. Student’s t-test, *** P <0.001. AA, amino acids; ST, *Salmonella*. **B)** WT cells were incubated with BafA1 and starved for AA for 3h in the presence or absence of ST. Fam134B puncta formation was observed by confocal microscopy. NR, Nutrient Rich; BafA1, Bafilomycin A1. Error bars indicate standard deviation. ANOVA, *** P <0.001. **C)** *FAM134B* KO cells were transfected with either FAM134B WT or FAM134B G216R and starved for AA for 6h or starved for 3h, followed by ST infection for 3h in AA starvation media. ER-phagy was measured by ss-RFP-GFP-KDEL processing. Actin was used as a loading control. **D)** Normalized Free RFP-Actin ratio from cells in C). Error bars indicate the standard deviation of 5 independent experiments. ANOVA, ***P <0.001; n.s, no significance. **E)** *FAM134B* KO were transfected with either FAM134B WT or G216R and starved for AA for 6h or starved for 3h followed by ST infection for 3h in AA starvation media. **F)** Average number of RFP puncta per cell from E) were quantified. ANOVA, ***P <0.001.

### The *Salmonella* effector SopF blocks ER-phagy

In order to promote invasion and replication, *Salmonella* expresses two type-III secretion systems, which deliver multiple bacterial effectors into the host cells^24^. We hypothesized that one of these effectors might be involved in *Salmonella* -mediated ER-phagy inhibition. To test our hypothesis, we repeated the ss-RFP-GFP-KDEL processing assay infecting them with different *Salmonella* effector mutants. Among the mutants tested, *Salmonella* defective for the phosphoinositide binding effector SopF showed the most complete and consistent loss of ER-phagy regulation. Analysis of ER-phagy by western blot and immunofluorescence in our reporter cells showed that SopF-deficient *Salmonella* was unable to supress ER-phagy compared to WT *Salmonella* infected controls under stimulated conditions (Fig. 4A-D). Together, these experiments indicate that SopF is necessary for *Salmonella* -induced ER-phagy repression. Next, we sought to determine if the expression of the SopF effector was sufficient to inhibit ER-phagy. To this end, ER-Phagy reporter cells were transfected with FLAG-SopF or control vector. We observed that SopF expression was sufficient to inhibit ER-phagy under stimulated conditions (Fig. 4E and 4F). Furthermore, SopF expression dramatically blocked endogenous FAM134B degradation by ER-phagy under stimulated conditions (Supp Fig. 3A). Altogether, these data indicate the bacterial effector SopF is required for *Salmonella* to inhibit ER-phagy.

**Fig. 4.**
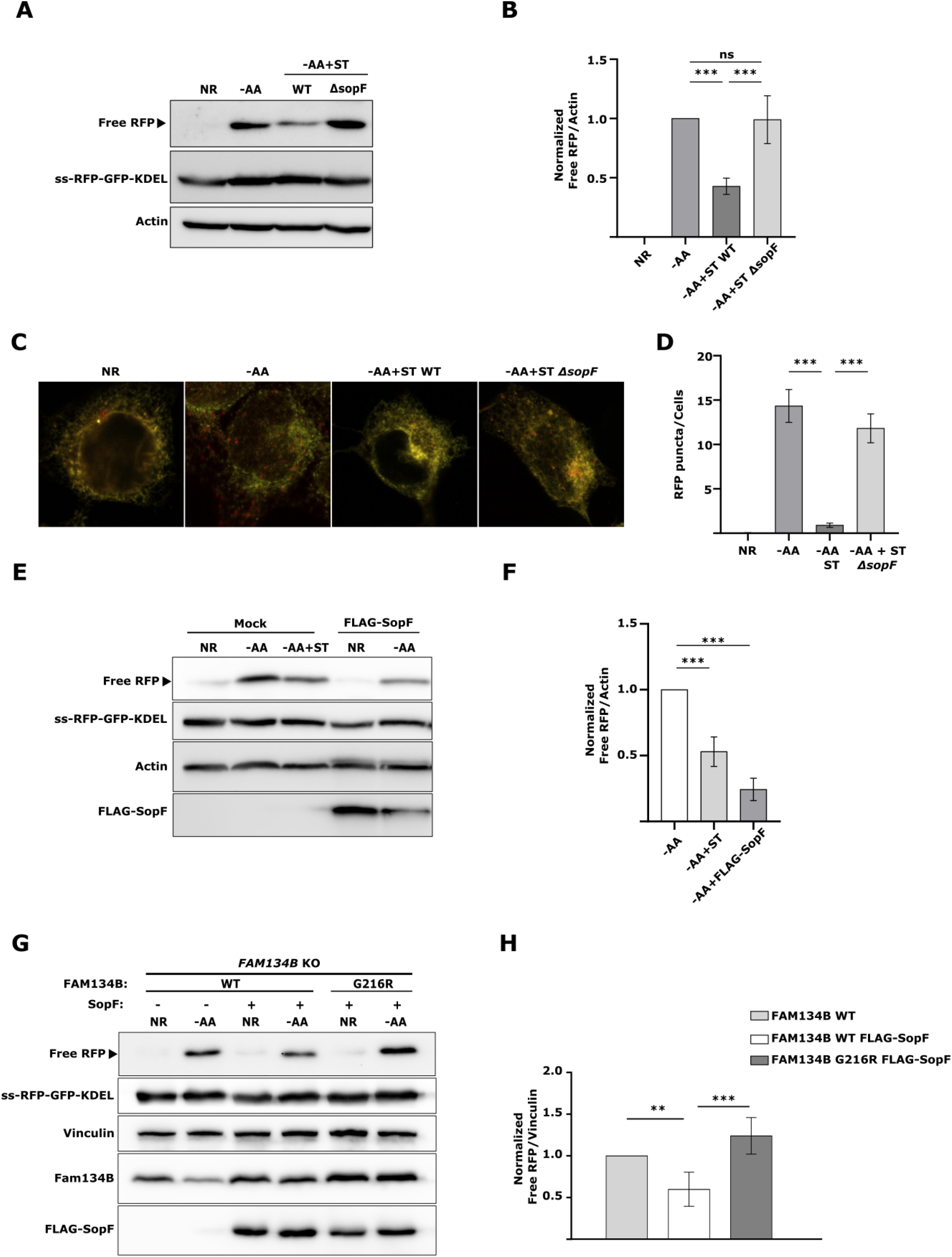
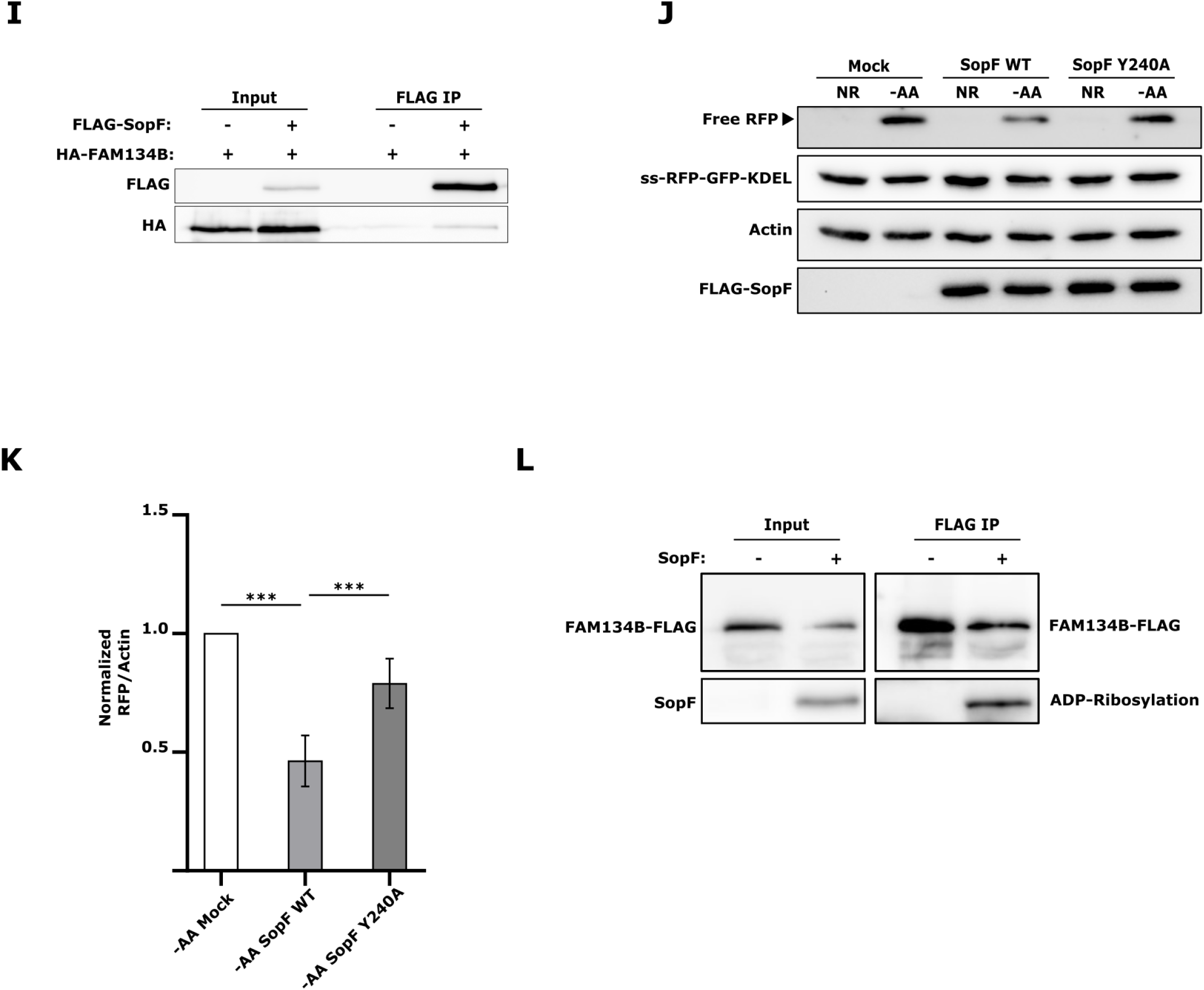
The *Salmonella* effector SopF blocks ER-phagy. **A)** WT HEK293 cells stably expressing the ss-RFP-GFP-KDEL reporter were starved for AA for 6h or starved for 3h followed by infection with ST WT or ST *ΔsopF* for 3h in AA starvation media. ER-phagy was measured by ss-RFP-GFP-KDEL processing. Actin was used as a loading control. NR, Nutrient Rich; AA, amino acids; ST, *Salmonella* Typhimurium. **B)** Normalized Free RFP-Actin ratio from cells in A). Error bars indicate the standard deviation of 4 independent experiments. ANOVA, ***P <0.001; n.s, no significance. **C)** WT cells were either AA starved for 6h or AA starved for 3h followed by ST WT or ST *ΔsopF* infection for 3h in starvation media, fixed and image by confocal microscopy. **D)** Average number of RFP puncta per cell from C) were quantified. ANOVA, ***P <0.001; n.s, no significance. **E)** WT cells transfected with either FLAG-SopF or a mock plasmid were starved for AA for 6h or starved for 3h followed by infection with ST WT for 3h in AA starvation media. ER-phagy was measured by ss-RFP-GFP-KDEL processing. Actin was used as a loading control. **F)** Normalized Free RFP-Actin ratio from cells in E). Error bars indicate the standard deviation of 3 independent experiments. ANOVA, ***P <0.001. **G)** *FAM134B* KO cells transfected with combinations of FAM134B WT, FAM134B G216R and FLAG-SopF were starved for AA for 6h. Vinculin was used as loading control. **H)** Normalized Free RFP-Vinculin ratio from cells in G). Error bars indicate the standard deviation of 5 independent experiments. ANOVA, **P <0.01; ***P <0.001. **I)** WT cells were transfected with HA-FAM134B and either FLAG-SopF or a mock plasmid. FLAG-SopF was immunoprecipitated. **J)** WT HEK293 cells transfected with either FLAG-SopF WT, FLAG-SopF Y240A or a mock plasmid were starved for AA for 6h ER-phagy was measured by ss-RFP-GFP-KDEL processing. Actin was used as a loading control. **K)** Normalized Free RFP-Actin ratio from cells in J). Error bars indicate the standard deviation of 6 independent experiments. ANOVA, ***P <0.001. **L)** *FAM134B* KO cells transfected with FAM134B-FLAG and either GFP-SopF or a mock plasmid and FLAG was immunoprecipitated. ADP-ribosylation was detected with a pan-ADP ribose binding reagent.

Because forcing FAM134B oligomerization with the G216R mutant could bypass *Salmonella* -mediated ER-phagy blockage, we sought to determine if SopF effects on ER-phagy could also be prevented by FAM134B G216R. Indeed, *FAM134B* KO cells co-expressing FLAG-SopF and FAM134B G216R were able to significantly induce ER-phagy when stimulated compared to cells transfected with FLAG-SopF and FAM134B WT (Fig. 4G and 4H). These data suggest *Salmonella* targets FAM134B oligomerization through SopF.

Recently, the crystal structure of SopF was solved revealing it to be a member of the ADP-ribosyltransferase superfamily. Indeed, SopF has been shown to catalyze the transfer of ADP-ribose from nicotinamide adenine dinucleotide (NAD^+^) to the v-ATPase subunit ATP6V0C leading to the inhibition of bacterial autophagy, but not general autophagy or other types of selective autophagy^25,26^. To test if SopF could directly interact with and target FAM134B, we co-immunoprecipitated FLAG-SopF and HA-FAM134B demonstrating a faint but reproducible interaction between both proteins (Fig. 4I). SopF mutations preventing its binding to the N-ribose (E325A) or the nicotinamide (Y224A, Y240A) group of NAD^+^ largely blocked SopF ADP-ribosylation activity^25^. Consistently, SopF Y240A failed to prevent ER-phagy activation as observed by a significant increase in RFP production compared to SopF WT. (Fig. 4J and 4K). Additionally, SopF mutants E325A and Y224A also showed increased ER-phagy compared to SopF WT (Supp Fig. 3B). Moreover, expression of SopF WT, but not the ADP-ribosylation mutant SopF Y240A, reduced the formation of FAM134B oligomers (Supp Fig. 3C). Finally, we used a pan-ADP-ribose binding reagent to test if SopF could ADP-ribosylate FAM134B. A clear band was observed when FAM134B was IP in the presence of SopF, which was absent in the control IP, suggesting SopF may directly ADP-ribosylate FAM134B (Fig. 4L). Altogether, our results indicate that SopF ADP-ribosylation activity is required for ER-phagy inhibition.

### FAM134B restricts *Salmonella* growth

To better understand the nature of FAM134B-mediated resistance to intracellular *Salmonella,* we next looked at the factors that could contribute to the *FAM134B* KO defect, such as bacteria vesicle escape, clearance, and growth. First, we infected WT and *FAM134B* KO cells with GFP *Salmonella* and quantified growth post-invasion for 4 hrs. We observed a similar level of bacterial internalization in both WT and *FAM134B* KO at 1 hr (Fig. 5A), indicating similar levels of infection rate. Interestingly, we observed a significant difference in growth rate between WT and *FAM134B* KO beginning at 2 hr post infection and continuing until the 4 hr endpoint. Escape from *Salmonella-* containing vesicles (SCV) is known to increase the growth rate of intracellular *Salmonella*, which is significantly higher in the cytosol. However, SCV escape in WT cells typically occurs well after 4 hrs post infection^27^ and results in the growth of rod-shaped *Salmonella* that are more spread out than those growing in the SCV. Given the early timepoint of divergence in growth rates and the morphology of internalized *Salmonella,* it is highly unlikely that *FAM134B* KO defects are a result of early escape from the SCV. We next sought to measure the impact of FAM134B on autophagic bacterial clearance, termed xenophagy. Autophagosomal degradation of *Salmonella* is mediated by LC3 recruitment to bacteria and autophagy-deficient cells have been shown to be more permissive for *Salmonella* growth^28^. However, the impact of ER-phagy in *Salmonella* clearance is unclear. Strikingly, LC3B-positive *Salmonella* showed a dramatic decrease in infected *FAM134B* KO compared to WT cells, suggesting a role for FAM134B and ER-phagy in *Salmonella* clearance (Fig. 5B and 5C). To determine if SopF is required for the suppression of LC3-positive puncta in *FAM134B* KO cells, we quantified the localization of LC3B to SopF-deficient *Salmonella.* We observed the previously reported increase of LC3B colocalization to *ΔsopF Salmonella* compared to WT *Salmonella* in our control cells^29^. However, SopF deletion exerted no significant difference in LC3B colocalization in *FAM134B* KO cells (Fig. 5D and 5E). Similarly, SopF overexpression significantly increased WT *Salmonella* levels in control cells, while overexpression of SopF in *FAM134B* KO cells showed no significant difference in *Salmonella* viability compared to mock transfected *FAM134B* KO cells (Fig. 5F), further indicating that SopF effects on *Salmonella* growth are linked to FAM134B. This indicates that the xenophagy defects in the *FAM134B* KO cells are presumably the cause of the increased bacterial susceptibility in these cells. Moreover, it reinforces our working model that SopF is inhibiting ER-phagy to promote intracellular *Salmonella* survival. The mechanisms that specifically link FAM134B-dependent ER-phagy to xenophagy are unclear at this time, but these results suggest an additional, yet undiscovered, pathway linking FAM134B to *Salmonella* restriction.

**Fig. 5.**
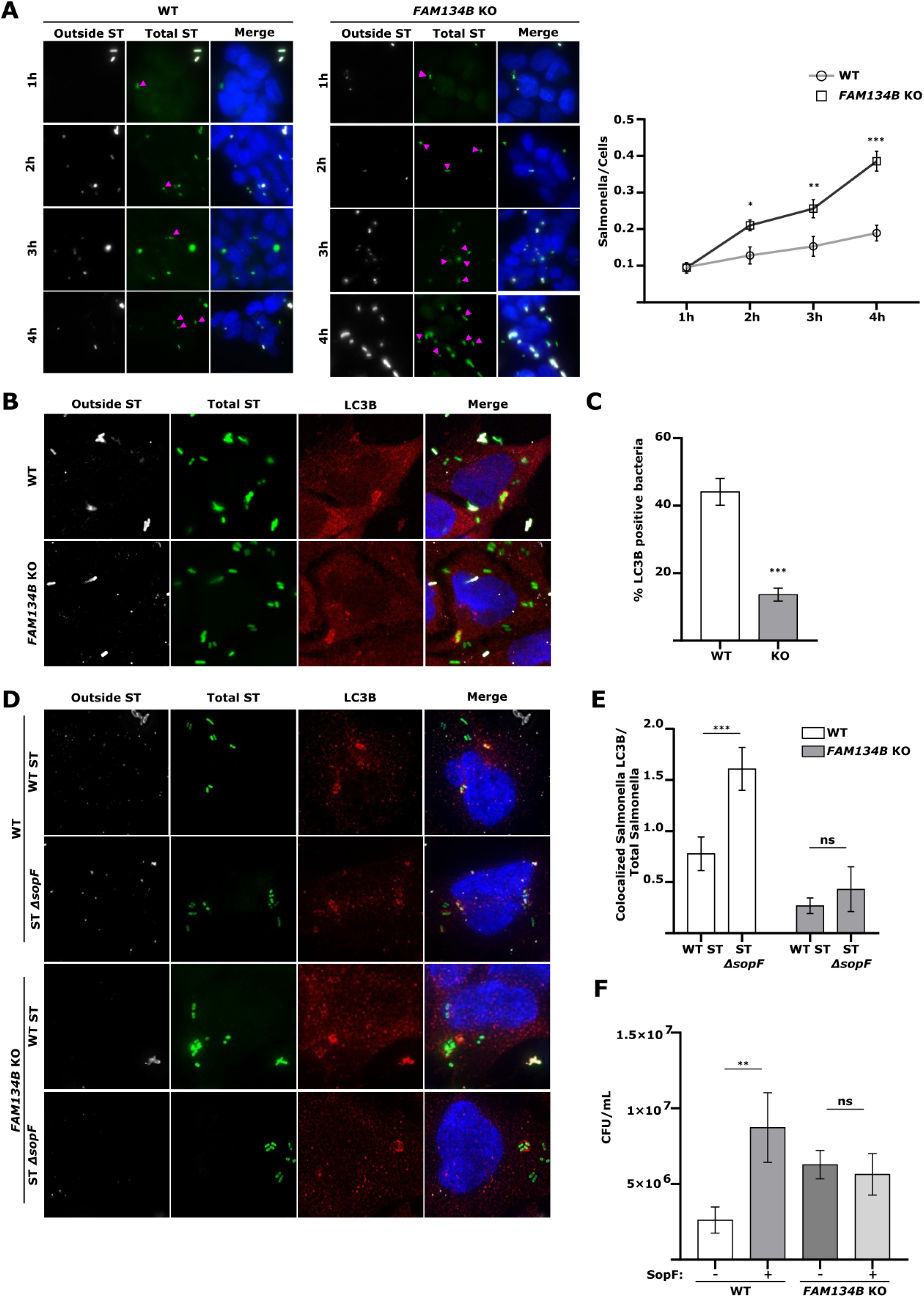
FAM134B restricts *Salmonella* growth. **A)** WT and *FAM134B* KO HEK293 cells were infected with ST for 1h followed by Gentamycin wash-off. Cells were fixed at indicated time points. ST, *Salmonella* Typhimurium. Error bars indicate standard deviation. ANOVA, *P <0.05; **P <0.01; ***P <0.001. **B)** WT and *FAM134B* KO cells were infected for 1h. Autophagic capture of ST was analyzed by immunostaining for LPS and LC3B**. C)** Percentage of ST colocalizing with LC3B in WT and *FAM134B* KO cells from B). Error bars indicate standard deviation. Student’s t test, ***P <0.001. **D)** WT and *FAM134B* KO cells were infected with either ST WT or ST *ΔsopF* for 1h. Autophagic capture of ST was analyzed by immunostaining for LPS and LC3B**. E)** The ratio between the number of ST colocalizing with LC3B and total internalized ST from D) is presented. Error bars indicate standard deviation. ANOVA, ***P <0.001; ns, no significance. **F)** WT and *FAM134B* KO were transfected with either FLAG-SopF or a mock plasmid followed by ST infection. Bacterial content was determined through the CFU. Error bars indicate standard deviation. ANOVA, **P <0.01.

### Infected *FAM134B* KO mice are more susceptible to *Salmonella* infection

We next sought to determine the pathophysiological effects of *Salmonella-* mediated inhibition of ER-phagy *in vivo*. To this end, we performed oral gavage *Salmonella* infection of WT and *FAM134B* KO mice^30^ and analyzed bacterial burden and intestinal damage 5 days post infection. Histochemical analysis of hematoxylin and eosin (H&E) stained intestine samples of infected WT mice presented mucosa and submucosa infiltration, along with occasional damage (Fig. 6A). However, infected *FAM134B* KO mice showed severe wall, mucosa and submucosa infiltration, as well as marked necrotic damage, fibrin formation and edema. Following H&E, mucosa and submucosa samples were stained to detect *Salmonella* levels, which showed a significant increase in bacterial burden in *FAM134B* KO compared to WT cells (Fig. 6B and 6C). These results are consistent with our *in vitro* data, highlighting FAM134B role in restricting *Salmonella* growth. Infections were repeated as described above and WT and *FAM134B* KO *Salmonella* load was measured by CFU in feces, spleen, and intestine. Consistent with immunofluorescence analysis, we observed significantly higher levels of *Salmonella* in the tissue and feces of *FAM134B* KO mice (Fig. 6D, 6E and 6F). We next tried to determine if our previous results extended to primary macrophages, key agents involved in the innate and adaptive immune response against *Salmonella* infection. Thus, we measured FAM134B-mediated ER membrane scission in starved bone marrow derived macrophages (BMDM) obtained from WT mice that were infected with either *Salmonella* WT or *ΔsopF*. As expected, ER-phagy activation induced FAM134B puncta formation, which was significantly blocked by *Salmonella* WT, but not SopF defective *Salmonella* (Fig. 6G and 6H), recapitulating our results in endothelial cells. Finally, BMDM obtained from *FAM134B* KO mice displayed significantly more *Salmonella* than WT BMDM after infection (Fig. 6I), further underlining the importance of FAM134B in anti-bacterial response in multiple tissues and cell types. Overall, our results indicate that FAM134B is an important factor in controlling bacterial infection *in vivo*.

**Fig. 6.**
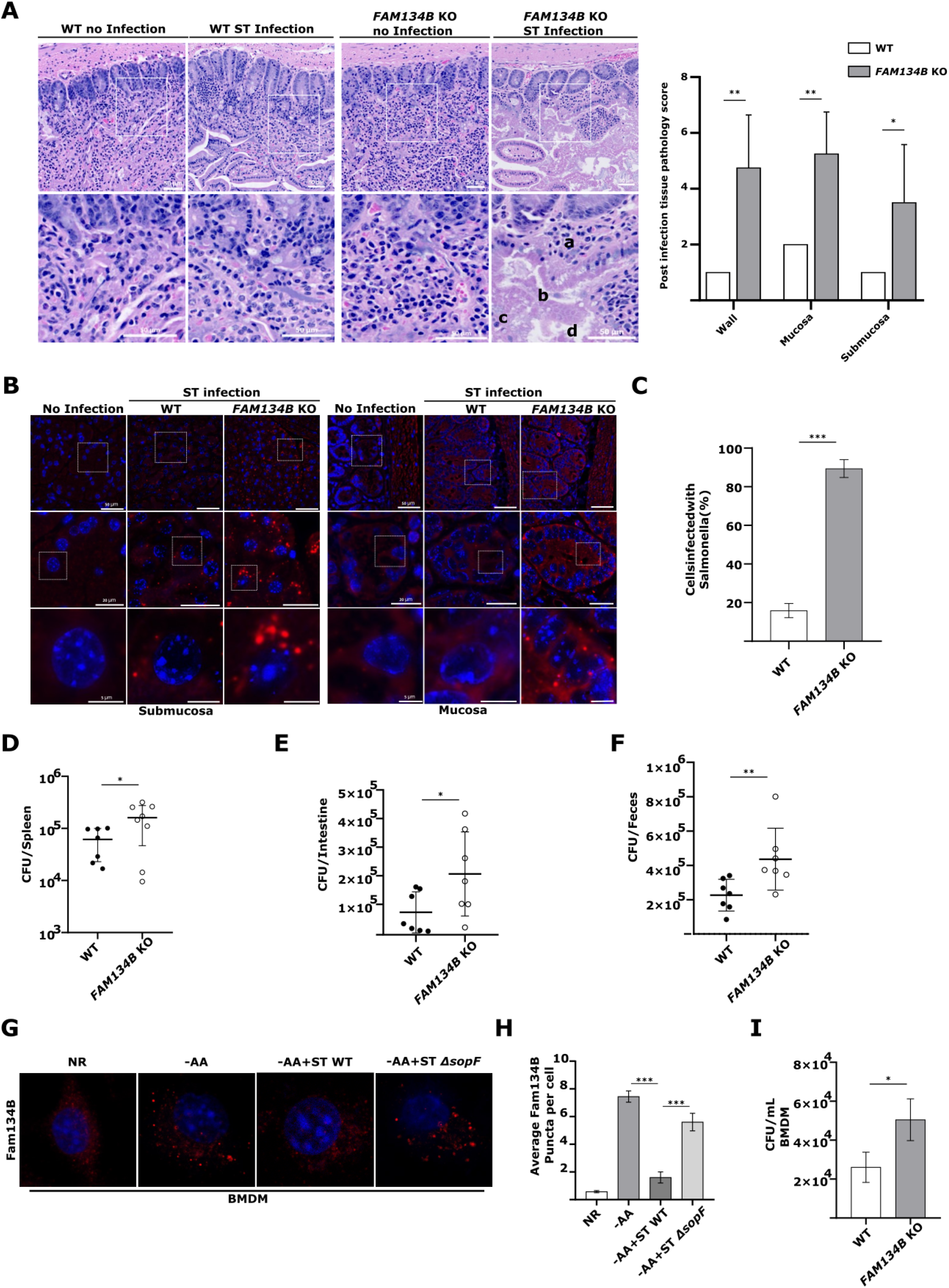
Infected *FAM134B* KO mice are more susceptible to *Salmonella* infection. **A)** WT and *FAM134B* KO mice were infected with ST through oral gavage and after 5 days their small intestine were fixed and stained with H&E. (a) infiltration, (b) necrosis, (c) fibrosis, and (d) edema. Pathology scores of post infected tissues are presented. Error bars indicate standard deviation. ANOVA, *P <0.05; **P <0.01. ST, *Salmonella* Typhimurium. **B)** WT and *FAM134B* KO mice intestine samples from A) were stained with DAPI and GFP to detect ST. **C)** The number of cells infected with ST from B) were quantified. Error bars indicate standard deviation. Student’s t test, ***P <0.001. **D, E and F)** WT and *FAM134B* KO mice were infected with ST through oral gavage and after 5 days their spleen, whole intestine and feces were collected. Bacterial content was determined through colony-forming unit (CFU). Error bars indicate standard deviation. Student’s t test, *P <0.05; **P <0.01. **G)** BMDM, bone marrow derived macrophages from WT mice were kept in NR media, AA starved for 6 hr or starve for 3 hr followed by ST WT or ST *ΔsopF* infection in starvation media for an additional 3 hr. All starve samples were treated with BaFA1. BMDM were then fixed and imaged by confocal microscopy. NR, Nutrient Rich; AA, amino acids, BafA1, Bafilomycin A1 **H)** Average number of FAM134B puncta per cell from G) were quantified. Error bars indicate standard deviation. ANOVA, ***P <0.001. **I)** BMDM from WT and *FAM134B* KO mice were infected with ST. Bacterial content was determined through colony-forming unit (CFU). Error bars indicate standard deviation. Student’s t test, *P <0.05.

## DISCUSSION

The ER is a dynamic and complex intracellular organelle with several critical functions involved in maintaining cell homeostasis and adapting to stress response^31^. The selective autophagic degradation of the ER, ER-phagy, aims to restore ER homeostasis through the degradation of portions of the ER and in turn regulate the size and morphology of the organelle. Recently, several studies have begun to uncover the link between ER-phagy and intracellular invasive pathogens. Here, we describe a novel mechanism by which the intracellular pathogen *Salmonella enterica* serovar Typhimurium specifically prevents the protein receptor FAM134B oligomerization in order to block ER-phagy and increase bacterial viability after infection.

We determined that *Salmonella* specifically targets FAM134B, but not TEX264, to block ER-phagy and increase bacterial viability, which was intrinsically linked to FAM134B ability to induce ER-phagy. A previous report highlighted how invasive bacteria capitalize on transforming the ER morphology to improve their viability. Upon *Legionella pneumophila* infection multiple ER regulatory proteins, including FAM134A, FAM134B, FAM134C, RTN4 and TEX264, are phosphoribosyl-linked ubiquitinated, triggering ER remodeling and membrane recruitment to bacterial containing vacuoles^18^. Further studies will be required to determine if other members of the FAM134 family isoforms or other proteins involved in the regulation of ER morphology and stability are also targeted by *Salmonella*.

We also identified that *Salmonella* -mediated ER-phagy blockage is linked to the bacterial effector SopF, specifically its ADP-ribosylation activity. Previously identified as an anti-bacterial autophagy inhibitor, SopF specifically disrupts the interaction between v-ATPase and ATG16L1 by ADP-ribosylating the v-ATPase subunit ATP6V0C^25,26^. In these experiments SopF was shown to inhibit anti-bacterial autophagy, but not canonical autophagy. However, ER-phagy was not tested^26^. Thus, it is likely that SopF can ADP-ribosylate multiple targets to both prevent *Salmonella* clearance and promote *Salmonella* growth. Interestingly, SopF-mediated ER-phagy blockage could be bypassed by expressing FAM134B G216R mutant, which promotes FAM134B oligomerization. Therefore, it is possible that SopF prevents FAM134B oligomerization by directly ADP-ribosylating FAM134B or indirectly by targeting other FAM134B interactors involved in the formation of multi-protein clusters required for ER-phagy^32^.

Finally, our *in vivo* results demonstrated the physiological importance of FAM134B in innate immunity. Infected *FAM134B* KO mice presented increased necrotic damage compared to infected WT mice. Additionally, *Salmonella* viability was increased in the spleen, intestine and feces of *FAM134B* deficient mice, as well as infected BMDMs, suggesting a yet uncharacterized but important role for FAM134B in the innate response against invading pathogens. Altogether, our data uncover a previously undiscovered mechanism by which *Salmonella* seeks to promote bacterial viability by targeting FAM134B-mediated ER-phagy and transform the host environment to suit their growth needs. We believe this study raises several important questions including the interplay between SopF targets in controlling *Salmonella* viability and the general ability of FAM134B-dependent restriction for other intracellular bacteria.

## MATERIAL AND METHODS

### Antibodies and reagents

HA-HRP (#Cat 2999), Fam134B (Cat# 83414), and phospho-S6K T389 (Cat#9234) antibodies were obtained from Cell Signaling Technology. Anti-LC3B (Cat#PM036 for immunofluorescence) antibodies were purchased from MBL. Pan-ADP-ribose binding reagent (MABE1016), Beta-actin (Cat#A5441 clone AC-15) and Vinculin (Cat#V9131) antibodies, as well as doxycycline hyclate (Cat#24390-14-5) were obtained from Sigma. DYKDDDDK Epitope Tag (Cat#NBP1-06712 for WB), anti-Tex264 (Cat#NBP1-89866) and LC3/MAP1LC3B (Cat#NB100-2220 for western blot) antibodies were purchased from Novus Biologicals. Anti-LPS FITC (Cat#sc-52223), antibodies were purchased from Santa Cruz Biotechnology. LPS (Cat#ab128709), and Anti-S6K (Cat#ab32529) were obtained from Abcam. Anti-RFP (Cat# 600-401-379) was obtained from Cedarlane. Anti-p62 (Cat#GP62-C) was purchased from Progen. Anti-Tex264 (Cat#25858-1-AP) from Proteintech. Bafilomycin A1 was obtained from Tocris (Cat#133410U). Digitonin (Cat#10188-874) was obtained from VWR.

### Cell culture

HEK293A were cultured in DMEM supplemented with 10% bovine calf serum (VWR Life Science Seradigm). Amino acid starvation medium was prepared based on the Gibco standard recipe, omitting all amino acids without the addition of non-essential amino acids and substitution with dialyzed FBS (Invitrogen). Doxycycline treatment of cells stably expressing ER-phagy probe was performed as previously described^7^.

### Transfection

All transfections were performed using polyethylenimine (PEI, medistore uOttawa). The samples were analyzed 48–72 h post-transfection.

### Generation of knock-out cell lines using CRISPR/Cas9

*FAM134B* and *TEX264* KO lines were generated in the HEK293A background utilizing CRISPR/Cas9 using primers that were previously described^7^. *FIP200* KO lines were generated using guide RNA sequence AGAGTGTGTACCTACAGTGC.

### Generation of stable cell lines

WT and KO clones were infected with lentiviruses carrying ss-RFP-GFP-KDEL as previously described^7^.

### Plasmids

Plasmids pEGFP-C1-SopF (#137734) and pMRX-INU-FLAG-FAM134B (#128260) were obtained from Addgene. SopF cDNA from pEGFP-C1-SopF was subcloned into FLAG-pcDNA to generate FLAG-SOPF plasmids. FAM134B cDNA from pMRX-INU-FLAG-FAM134B was subcloned into HA-pcDNA and c-FLAG-pcDNA to generate HA-FAM134B and FAM134B-FLAG plasmids, respectively. All constructs were generated using fast-cloning as previously described^33^.

### Site-directed mutagenesis

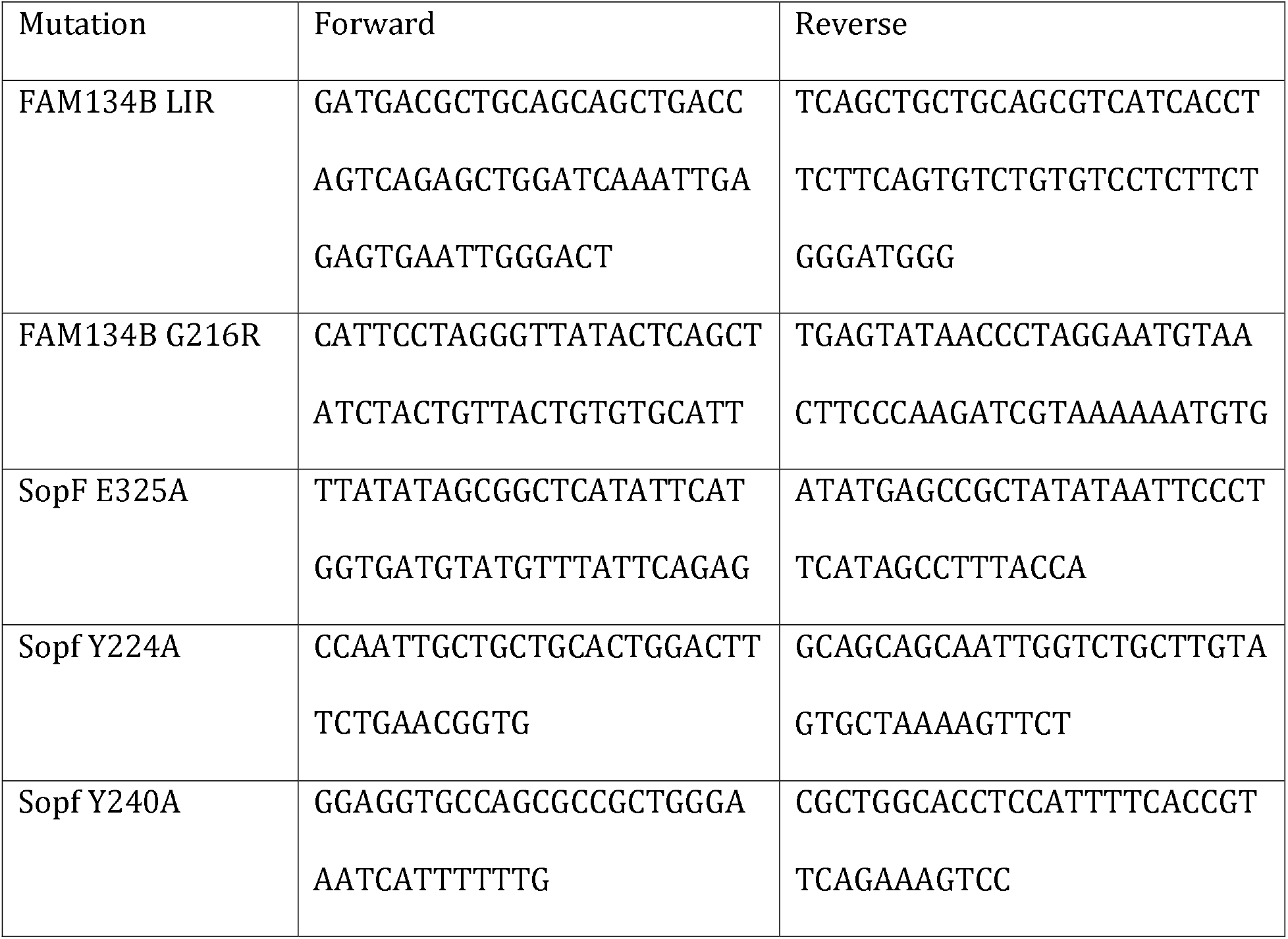
Primers used for making FAM134B LIR mutation.

Site-directed mutagenesis was performed as previously described^34^. Specificity of mutagenesis was analyzed by direct sequencing.

### Bacterial strains

Wild-type (SL1344) *Salmonella* strain was a gift from Dr. Subash Sad (University of Ottawa). *Salmonella ΔsopF* strain (SL1344) was a gift from Dr. John Brumell. Bacteria were grown in Luria-Bertani broth (Fisher).

### Bacterial infection

*Salmonella* was grown in 4 ml of LB broth at 37°C at 250 rpm. Overnight cultures of *Salmonella* were diluted 30-fold and allowed to grow until reaching an OD_600_ of 1.5, followed by centrifugation of 10,000 *g* for 2 min, and the resulting pellet was resuspended in 1 ml of phosphate-buffered saline (PBS). Bacterial stock was then diluted to multiplicity of infection (MOI) of 180 in DMEM supplied with 10% heat-inactivated bovine calf serum for infection or amino acid starvation media. Cells cultured in antibiotic-free medium were infected with *Salmonella* infection and maintained at 37°C in a 5% CO2 environment for the specified duration. Prior to analysis, cells were washed once with PBS and lysed directly using 1x denaturing SDS sample buffer.

### Western blot and immunoprecipitation

Whole-cell lysates were prepared by direct lysis with 1x SDS sample buffer, followed by boiling for 10 minutes at 95°C and resolved by SDS–PAGE. Immunoprecipitation cells were harvested in mild lysis buffer (MLB) containing 10 mM Tris pH 7.5, 10 mM EDTA, 100 mM NaCl, 50 mM NaF, and 1% NP-40, supplemented with protease and phosphatase inhibitor cocktails (including EDTA from APExBIO). The lysates were then centrifuged at maximum speed for 10 minutes to remove cell debris. Anti-FLAG affinity gel beads (Sigma) were washed once with MLB and then incubated with cell lysates for 1.5 hours, followed by a single wash with MLB containing inhibitors and four quick washes with MLB alone. The beads were subsequently boiled in a 1x denaturing sample buffer for 10 minutes before being resolved by SDS–PAGE. Imaging was conducted using the ChemiDoc™ Touch imaging system (Bio-Rad).

### Immunofluorescence

Cells were seeded onto IBDI-treated coverslips and allowed to adhere overnight. Following treatments, cells were fixed in 4% paraformaldehyde in PBS for 15 minutes and then permeabilized with 50 μg/ml digitonin in PBS for 10 minutes at room temperature. Subsequently, cells were blocked using a blocking buffer (1% BSA and 2% serum in PBS) for 45 minutes, followed by incubation with primary antibodies in the same buffer for 1 hour at room temperature. After incubation, samples were washed three times in PBS and once in blocking buffer before being incubated in secondary antibodies for 1 hour at room temperature. Slides were washed three times in PBS, stained with DAPI, and mounted. Imaging was conducted using an inverted epifluorescent Zeiss AxioObserver.Z1 microscope. For staining bacterial localization (inside/outside), cells were first incubated with an anti-LPS antibody for 1hr and followed by secondary antibody incubation for 1hr in a blocking buffer before permeabilization. This was followed by three PBS washes between steps. Epifluorescent microscopy images were analyzed using an automated protocol implemented in ImageJ software to minimize bias. The same protocol was consistently applied to each field of view and across all samples. An average of seven unique fields of view from representative experiments were selected for quantification.

### Cyto-ID Autophagy Detection Kit assay

Cells were seeded onto ibidi 8 well µ-Slides (Ibidi, cat. 80826) and allowed to adhere overnight. Subsequently, cells were subjected to amino acid starvation with or without *Salmonella* infection. Following treatment, cells were incubated in DMEM without phenol red containing Cyto-ID autophagy detection stain (Enzo, ENZ-KIT175-0050) for 30 minutes, then washed with PBS and re-incubated with either complete DMEM without phenol red or amino acid media. Images were acquired and deconvolved using an inverted epifluorescent Zeiss AxioObserver.Z1 microscope.

### Colony-forming unit assay

Cells were infected with *Salmonella* at a multiplicity of infection (MOI) of 180 for 1 hour. Subsequently, the infected cells were rinsed three times and treated with media containing 100 μg/ml Gentamicin for 0.5 hours, followed by a 4-hour incubation with media containing 50 μg/ml Gentamicin. After incubation, the samples were washed three times with PBS and then lysed using CFU buffer (0.1% Triton X-100 and 0.01% SDS in PBS). The lysates obtained were subjected to serial dilution (1:50, 1:75, and 1:100) and plated onto LB agar plates containing Streptomycin. The plates were incubated at 37°C for 16–18 h, and the colonies were counted to determine the number of CFU.

### Immunohistochemistry staining

Samples were rinsed three times with PBS, treated with 3% H_2_O_2_ (in PBS) for 10 min, and washed three times with PBS. Blocked with protein block serum-free (catalog no. X0909 Dako) for 2 h, stained with primary antibody overnight at 4 °C (GFP 1:150 catalog no. Sigma #G1544), washed three times with PBS, incubated with secondary antibody (Alexa Fluor 555 anti-rabbit, catalog no. A31572, 1:1,000) for 111h, washed once with PBS, stained with DAPI (211mg11ml−1, Roche Diagnostics) for 10 min, washed three times with PBS and cover slip-mounted with Fluoromount-G mounting solution (Invitrogen, 00-4958-02). All treatments were done at room temperature unless otherwise stated. Images were acquired using a Zeiss LSM 800 AxioObserver Z1 Confocal Microscope

### *In vivo* experiments

WT and *FAM134B* KO mice were fasted for 3 hours prior to treatment with streptomycin (200 mg/mL). Food and water were replaced in the cage 3 hours post-treatment. The following day, mice were fasted for 3 hours prior to infection with ST-GFP (1x108 CFU) through oral gavage. Food and water were replaced in the cage 3 hours post-infection. 5 days post infection post-infection. 5 days post-infection, intestines were harvested and fixed in 10% formalin for 2 days. Then rinsed 3 times with 30% sucrose 3 times. The samples were then dehydrated and paraffin-embedded and paraffin-embeddedparaffin-embedded, sectioned into 4 µm thick slices and mounted onto glass microscope slides. Prior to staining, samples were rehydrated and deparaffinized. Antigen-retrieval for both groups was performed in pH 9.0 EDTA solution, at 110°C for 12 minutes in a microwave processor (Histo5, Milestone). For CFU assays after 5 days post-infection, the spleen, intestine, and colon of infected mice were harvested for quantification of the bacterial burden on selection agar plates.

### Statistical analysis

Error bars for western blot analysis represent the standard deviation between densitometry data collected using ImageJ software from at least three unique biological experiments. Statistical analyses were performed using GraphPad Prism 8. Statistical significance was determined using either Student’s t-test or ANOVA. Differences with a P value <0.05 or lower were considered significant. significant. *p<0.05, **p<0.01, ***p<0.001. The number of independent experiments (n), statistical measurements tests utilized, dispersion of measurements, and significance are described in the figure legends.

### Ethics compliance

All experiments, including animals, were approved by the University of Ottawa Animal Care Committee and are in accordance with the Guidelines of the Canadian Council on Animal Care.

## Abbreviations

Atg: autophagy-related
WT: wild-type
KO: knock-out
GFP: Green Fluorescent Protein
RFP: Red Fluorescent Protein
ss: signal sequence
BAFA1: Bafilomycin A1
CFU: Colony-Forming Unit
LIR: LC3-interacting region
NAD+: nicotinamide adenine dinucleotide
SCV: Salmonella containing vesicles
H&E: hematoxylin and eosin
BMDM: Bone Marrow Derived Macrophages

## Acknowledgements

We would like to thank Dr. John Brumell for sharing *Salmonella* mutants. We would also like to thank Karyn King for providing *FIP200* KO cell lines and Dr. Rudolf Mueller for scoring H&E staining. A big thank you to Zaida Ticas, Marjan Khalili and Mufida Alazzabi from the Louise Pelletier HCF at the University of Ottawa for all their help.

The authors would like to acknowledge the assistance of StemCore Laboratories Genomics Core Facility (OHRI, uOttawa), RRID:SCR_012601. The authors must acknowledge the Cell Biology and Image Acquisition Core (RRID: SCR_021845) funded by the University of Ottawa, Ottawa, Natural Sciences and Engineering Research Council of Canada, and the Canada Foundation for Innovation. We gratefully acknowledge the IHC tissue samples processing services provided by the Louise Pelletier HCF (RRID: SCR_021737) at the University of Ottawa.

This work was supported by the Canadian Institutes of Health Research (CIHR), funding reference number 181799 (D.G) & 376756 (R.C.R), 153034 (R.C.R), and Natural Sciences and Engineering Research Council of Canada #2023-05587 (R.C.R). R.M.A received support from the Ottawa Institute of System Biology and the Centre for Infection, Immunity and Inflammation.

## Author Contributions

D.G., R.M.A. and R.R. designed *in vitro* experiments. D.G. and R.M.A. performed *in vitro* experiments. D.G., R.M.A., R.EH., S.S. and R.R. designed mice experiments. D.G., R.M.A. and R.EH. performed mice experiments. R.M scored H&E samples. D.G., R.M.A. and R.R. wrote the manuscript. M.M provided *FAM134B* KO mice. All authors discussed the results and commented on the manuscript.

## Figure Legends

**Supplementary Fig. 1. Related to Figure 1. *Salmonella* infection does not affect lysosomal activity. A)** ER-phagy measurement example by ss-RFP-GFP-KDEL processing. WT and *FIP200* KO HEK293 cells stably expressing the ss-RFP-GFP-KDEL reporter were either starved for AA or treated with Torin 1 for 6h. ER-phagy was measured by ss-RFP-GFP-KDEL processing. Actin was used as a loading control. AA, Amino Acids; S.E, Short Exposure; L.E, Long Exposure. **B)** WT HEK293 cells were either kept in NR, starved for AA in the absence or presence of ST or infected with ST in NR conditions. Non-selective autophagy was measured by CytoID. NR, Nutrient Rich; ST, *Salmonella* Typhimurium. **C)** WT and *FIP200* KO cells stably expressing the ss-RFP-GFP-KDEL reporter were starved for AA for 6h in the presence or absence of BafA1 or starved for 3h followed by ST infection for 3h in AA starvation media with or without BafA1. ER-phagy was measured by ss-RFP-GFP-KDEL processing. Non-selective autophagy activity was measured by p62/SQSTM degradation. Actin was used as a loading control. BafA1, Bafilomycin A1.

**Supplementary Fig. 2. Related to Figure 2. *FAM134B*, *TEX264* KO and Additional CFU. A)** Fam134B, Tex264 and Actin levels were determined by western blot in HEK293 WT, *FAM134B* KO and *TEX264* KO cells **B)** WT and *FAM134B* KO cells transfected with either FAM134B WT, FAM134B LIR mutant or a mock plasmid were infected with ST. Bacterial content was determined through colony-forming unit (CFU). Error bars indicate standard deviation. ANOVA, *P <0.05; ***P <0.001. ST, *Salmonella* Typhimurium.

**Supplementary Fig. 3. Related to Figure 4. SopF ADP-ribosylation mutants fail to block ER-phagy. A)** WT HEK293 cells expressing FLAG-SopF or a mock plasmid were AA starved for 2h. Fam134B, Actin and FLAG-SopF levels were determined by western blot. NR, nutrient rich; AA, amino acids. **B)** WT cells transfected with either FLAG-SopF WT, FLAG-SopF WT E325A, FLAG-SopF Y224A, Y240A or a mock plasmid were starved for AA for 6h. ER-phagy was measured by ss-RFP-GFP-KDEL processing. Actin was used as a loading control. **C)** *FAM134B* KO cells transfected with FAM134B-FLAG and either GFP-SopF WT, GFP-SopF Y240A or a mock plasmid were starved for AA for 3h. FLAG was immunoprecipitated.

