## Supplementary figures and images for "The ER-phagy receptor FAM134B is targeted by *Salmonella* Typhimurium to promote infection"

### Supplemental Figure 1

#### Supp Fig.1 Related to Fig1

A

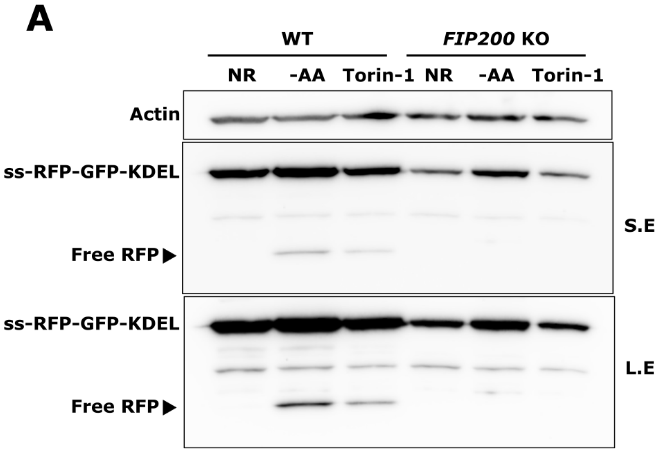


B

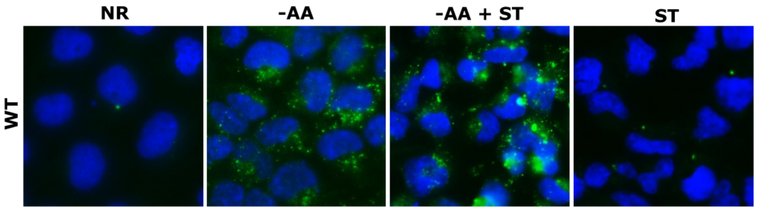


C

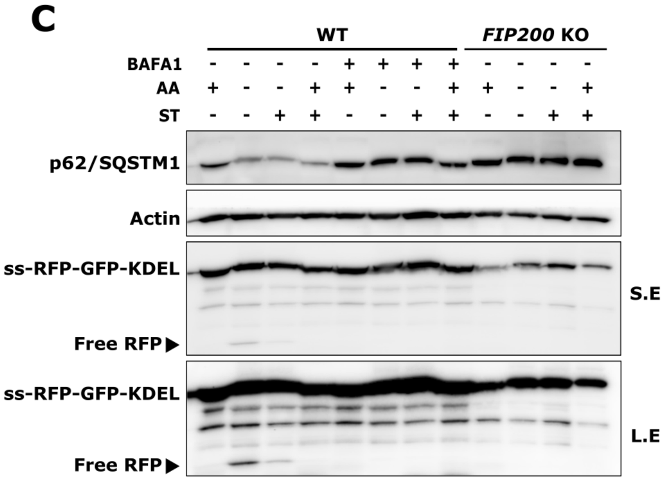

### Supplemental Figure 2

**Supp Fig. 2 Related to Fig2**

**A**

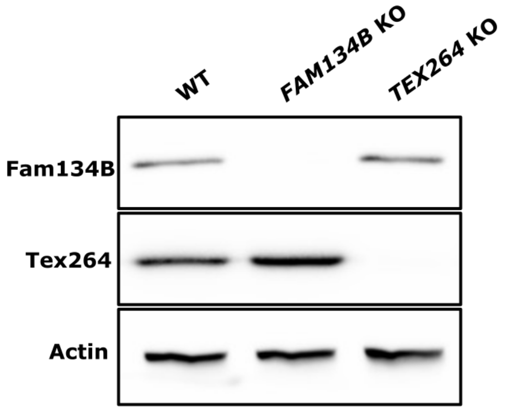


**B**

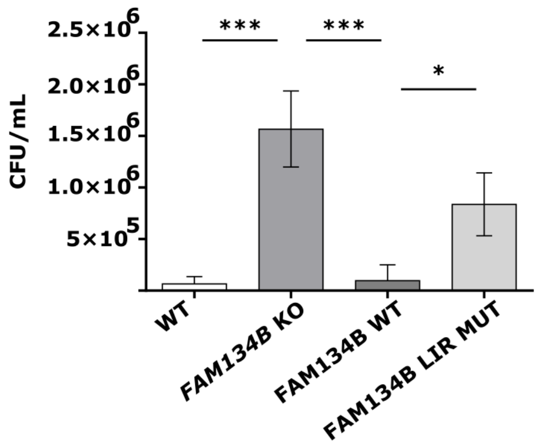

### Supplemental Figure 3

**Supp Fig. 3 Related to Fig4**

**A**

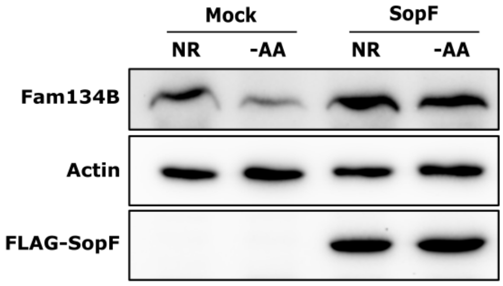


**B**

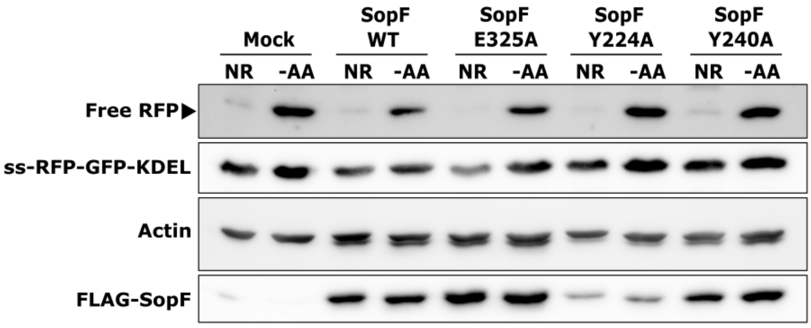


**C**

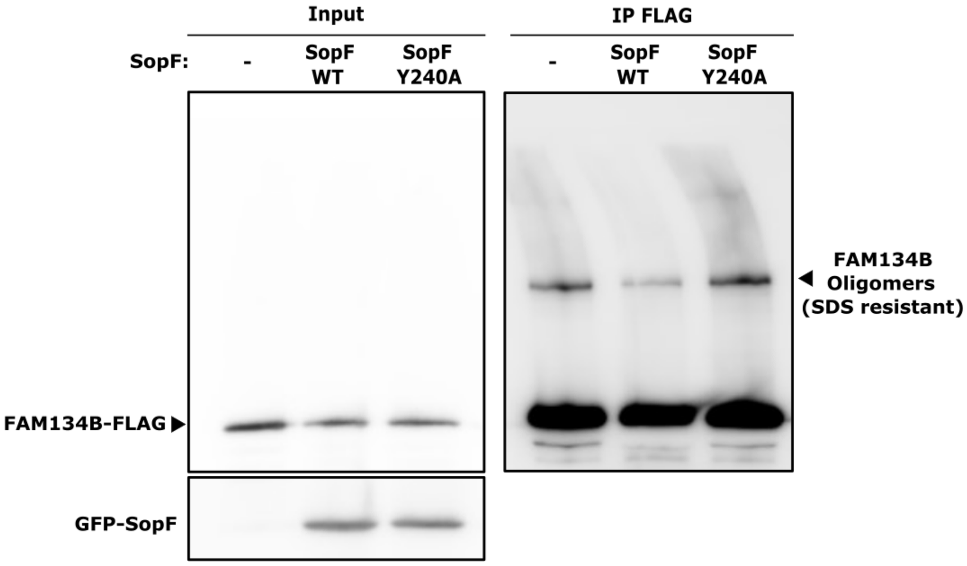
